# Impact of physico-chemical parameters on the denitrification and the co-occurrence of *Methylophaga nitratireducenticrescens* and *Hyphomicrobium nitrativorans* in a methanol-fed marine denitrifying biofilm

**DOI:** 10.1101/602896

**Authors:** Geneviève Payette, Valérie Geoffroy, Christine Martineau, Richard Villemur

## Abstract

**Background:** The biofilm of a continuous, methanol-fed, fluidized denitrification system that treated a marine effluent is composed of multi-species microorganisms, among which *Hyphomicrobium nitrativorans* NL23 and *Methylophaga nitratireducenticrescens* JAM1 are the principal bacteria involved in the denitrifying activities. This biofilm can be cultured at laboratory-scale under batch mode conditions without losing the denitrifying activities. Here, we report the capacity of the denitrifying biofilm to sustain changes to specific physico-chemical parameters, and the impact of these changes on the denitrification performance and the co-occurrence of *H. nitrativorans* and *M. nitratireducenticrescens*.

**Methods:** The original biofilm (OB) taken from the denitrification system was acclimated to an artificial seawater (ASW) medium under anoxic conditions to generate the Reference biofilm cultures (300 mg-NO_3_^−^-N/L, 23°C). In the first set of assays, the Reference biofilm cultures were subjected to short exposures (1-3 days) of a range of NaCl, methanol, nitrate (NO_3_^−^) and nitrite (NO_2_^−^) concentrations, and to different pHs and temperatures. In the second set of assays, the OB was acclimated in an ASW-modified medium for five weeks i) to a range of NaCl concentrations (0% to 8%), ii) to four combinations of NO_3_^−^ concentrations and temperatures, iii) to NO_2_^−^, and iv) under oxic conditions. Finally, the OB was acclimated to the commercial Instant Ocean (IO) medium. The growth of the biofilm and the dynamics of NO_3_^−^ and NO_2_^−^ were determined. The levels of *M. nitratireducenticrescens* and *H. nitrativorans* were measured by qPCR in these cultures.

**Results:** The denitrifying capacities of the OB was preserved in the Reference biofilm cultures. In these cultures, however, *H. nitrativorans* NL23 decreased by three orders of magnitude in concentration with the occurrence of the same magnitude of a new denitrifying bacterial strain (*M. nitratireducenticrescens* GP59). Results from the first set of assays showed that the Reference biofilm cultures can sustain denitrifying activities in most of the tested conditions. Inhibition occurred when these biofilm cultures were exposed at pH 10 or with 1.5% methanol. Results from the second set of assays showed the persistence of *H. nitrativorans* NL23 in the biofilm cultures acclimated to low NaCl concentrations (0% to 1.0%). Poor biofilm development occurred in biofilm cultures acclimated to 5% and 8% NaCl. Finally, high proportion of *H. nitrativorans* NL23 was found in the IO biofilm cultures.

**Conclusions:** These results confirm the plasticity of the marine methylotrophic denitrifying biofilm in adapting to different conditions. The NaCl concentration is a crucial factor in the dynamics *of H. nitrativorans* NL23, for which growth was impaired above 1% NaCl in the ASW-based biofilm cultures in favor of *M. nitratireducenticrescens* GP59. This study contributes to the understanding on the population dynamics of co-occurring bacteria performing denitrifying activities in biofilm under seawater environment. This could benefit in the development of optimal denitrifying bioprocess under marine conditions.

## Introduction

Denitrification takes place in bacterial cells where N oxides serve as terminal electron acceptor instead of oxygen (O_2_) for energy production when oxygen depletion occurs, leading to the production of gaseous nitrogen (N_2_). Four sequential reactions are strictly required for the reduction of NO_3_^−^ to gaseous nitrogen, via nitrite (NO_2_^−^), nitric oxide and nitrous oxide, and each of these reactions is catalyzed by different enzymes, namely NO_3_^−^ reductases (Nar and Nap), NO_2_^−^ reductases (Nir), nitric oxide reductases (Nor) and nitrous oxide reductases (Nos) (Philippot and Hojberg, 1999; Richardson *et al.*, 2001; Kraft *et al.*, 2011). These biological activities have been used with success to remove NO_3_^−^ in different wastewater treatment processes (Tchobanoglous *et al.*, 2003). The Montreal Biodome, a natural science museum, operated a continuous, fluidized methanol-fed denitrification reactor to remove NO_3_^−^ that accumulated in the 3 million-L seawater aquarium. The fluidized carriers in the denitrification reactor were colonized by naturally occurring multispecies bacteria to generate a marine methylotrophic denitrifying biofilm to be composed by around 15-20 bacterial species (Labbé *et al.*, 2003). Fluorescence in situ hybridization experiments on the biofilm showed that the methylotrophic bacteria *Methylophaga* spp. and *Hyphomicrobium* spp. account for 60 to 80% of the bacterial community (Labbé *et al.*, 2007).

*Hyphomicrobium* spp. are methylotrophic bacteria that are ubiquitous in the environment (Gliesche *et al.*, 2005). They have also been found in significant levels in several methanol-fed denitrification systems treating municipal or industrial wastewaters or a seawater aquarium, and they occurred often with other denitrifying bacteria such as *Paracoccus* spp., *Methylophilales* or *Methyloversatilis* (CatalanSakairi *et al.*, 1997; Lemmer *et al.*, 1997; Layton *et al.*, 2000; Ginige *et al.*, 2004; Baytshtok *et al.*, 2008; Wang *et al.*, 2014; Rissanen *et al.*, 2017). Their presence correlates with optimal performance of bioprocess denitrifying activities. *Methylophaga* spp. are methylotrophic bacteria isolated from saline environments (Bowman, 2005; Boden, 2012). They have been found in association with diatoms, phytoplankton blooms, marine algae, which are known to generate C1 carbons (Li *et al.*, 2007; Bertrand *et al.*, 2015; Landa *et al.*, 2018).

Two bacterial strains representatives of *Methylophaga* spp. and *Hyphomicrobium* spp. were isolated from the Biodome denitrifying biofilm. The first one, *Methylophaga nitratireducenticrescens* JAM1, is capable of growing in pure culture under anoxic conditions by reducing NO_3_^−^ in NO_2_^−^, which accumulates in the medium (Auclair *et al.*, 2010; Villeneuve *et al.*, 2013). It was later shown to be able to reduce NO and N_2_O to N_2_ (Mauffrey *et al.*, 2017). The second bacterium, *Hyphomicrobium nitrativorans* NL23, is capable of complete denitrification from NO_3_^−^ to N_2_ (Martineau *et al.*, 2013b). These two strains are the main bacteria responsible of the dynamics of denitrification in the biofilm. The genomes of *M. nitratireducenticrescens* JAM1 and *H. nitrativorans* NL23 were sequenced (Villeneuve *et al.*, 2012; Martineau *et al.*, 2013a). Strain JAM1 contains two Nar-type NO_3_^−^ reductase gene clusters (Nar1 and Nar2). Both Nar systems contribute to the reduction of NO_3_^−^ to NO_2_^−^ during strain JAM1 growth (Mauffrey *et al.*, 2015). Gene clusters encoding two cNor and one Nos systems are present and expressed in strain JAM1. Their presence correlates with the reduction of NO and N_2_O by stain JAM1 (Mauffrey *et al.*, 2017). A dissimilatory NO-forming NO_2_^−^ reductase gene (*nirS* or *nirK*) is absent, which correlates with accumulation of NO_2_^−^ in the culture medium during NO_3_^−^ reduction. The genome of *H. nitrativorans* NL23 contains the operons that encode for the four different nitrogen oxide reductases, among which a Nap-type NO_3_^−^ reductase (Martineau *et al.*, 2015).

We have initiated a study to assess the performance of the Biodome denitrifying biofilm subjected to different conditions. In the first part of this study reported by Geoffroy *et al.* (2018), the original biofilm (OB) taken from the Biodome denitrification system was cultured under batch-mode, anoxic conditions at laboratory scale to an artificial seawater (ASW) instead of the commercial Instant Ocean (IO) used by the Biodome. We noticed in the ASW-acclimated biofilm cultures important decreases in the concentration of *H. nitrativorans* NL23 without the loss of denitrifying activities and the occurrence of a new denitrifying bacterial strain, *M. nitratireducenticrescens* GP59. The genome of strain GP59 is highly similar of that of strain JAM1, but contains a *nirK* gene, which is missing in strain JAM1 (Geoffroy *et al.*, 2018).

**Table 1:**
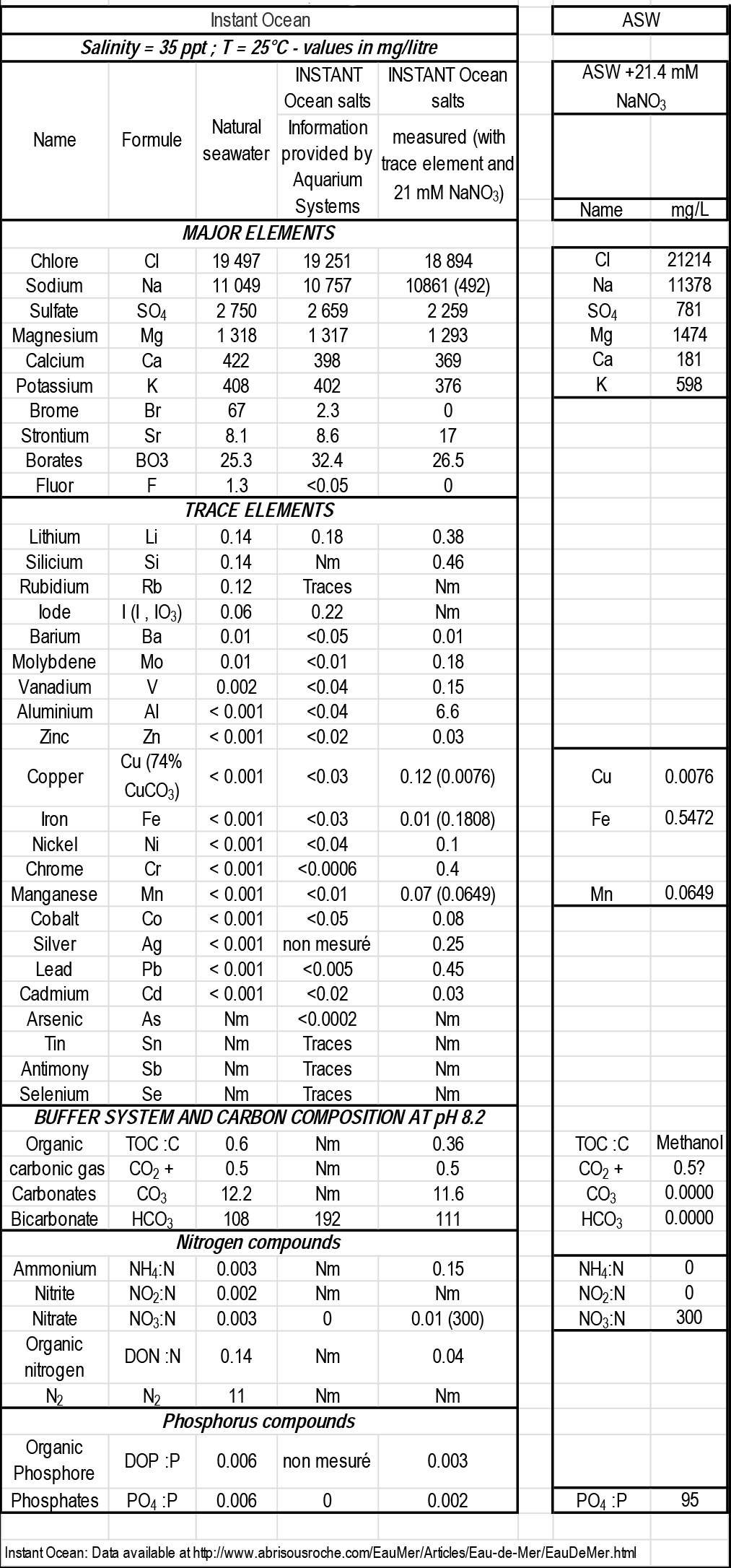
Composition of natural seawater, the INSTANT OCEAN brand salt and the artificial seaeater (ASW) used in this study.

In the second part of this study, which is reported here, further experiments were performed to assess the impact of changing specific physico-chemical parameters on the denitrifying biofilm. These parameters include varying the concentrations of NaCl, methanol, NO_3_^−^ and NO_2_^−^, varying the pH and temperature, and finally the type of seawater. The impact of these changes on the denitrifying activities and on the population dynamics of *M. nitratireducenticrescens* JAM1 and GP59 and *H. nitrativorans* NL23 was then measured. Our results revealed that the biofilm can sustain denitrifying activities in most of the tested conditions. It also showed a strong impact of the NaCl concentrations and the type of seawater on the persistence of strain NL23 in the denitrifying biofilm. To our knowledge, this is the first systematic study on the co-evolution of *Methylophaga* and *Hyphomicrobium* strains in a marine biofilm system.

## Material and Methods

### Seawater media

The artificial seawater (ASW) medium was composed of (for 1 liter solution): 27.5 g NaCl, 10.68 g MgCl_2_*6H_2_O, 2 g MgSO_4_*7H_2_O, 1 g KCl, 0.5 g CaCl_2_, 456 μL of FeSO_4_*7H_2_O 4 g/L, 5 mL of KH_2_PO_4_51,2 g/L, 5 mL of Na_2_HPO_4_34 g/L. The Instant Ocean (IO) seawater medium was bought from Aquarium systems (Mentor, OH, USA) and dissolved at 30 g/L. Estimation of its composition and comparison with the ASW medium is provided in Table 1. Both media were supplemented with 1 mL of trace element (FeSO_4_*7H_2_O 0.9 g/L, CuSO_4_*5H_2_O 0.03 g/L and MnSO_4_*H_2_O 0.2 g/L) and with NaNO_3_ (Fisher Scientific Canada, Ottawa, ON, Canada) at various concentrations, depending on the experiment. The pH was adjusted (NaOH) at 8.0 before autoclaving. Filtered-sterilize methanol (Fisher Scientific) was then added as a carbon source to support bacterial growth, also at various concentrations depending on the experiment. The sterile media were distributed (60 ml) in sterile serologic vials, which included 20 « Bioflow 9 mm » carriers (Rauschert, Steinwiessen, Allemagne). Prior to use, these carriers were washed with HCl 10% (v/v) for 3 hrs, rinsed with water and autoclaved. The vials were purged of oxygen for 10 min with nitrogen gas (Praxair, Mississauga, ON, Canada) and sealed with sterile septum caps.

**Figure 1.**
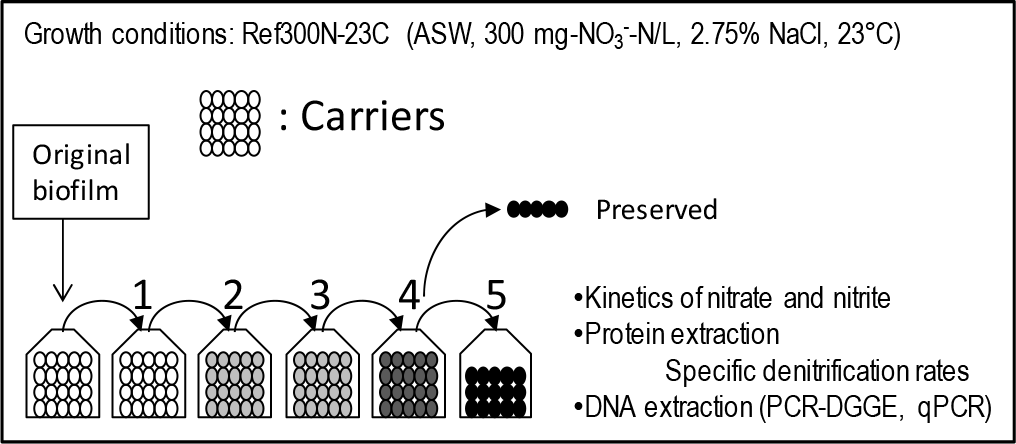
Development of the Reference biofilm cultures (Ref300N-23C) The original biofilm was thawed, scrapped from the carriers and distributed to vials containing the ASW medium supplemented with 300 mg NaNO_3_-N/L and 0.15% methanol and 20 free carriers. These carriers were transferred 5 times in fresh medium. Concentrations of NO_3_^−^ and NO_2_^−^ were measured at regular intervals. Methanol and NaNO_3_ were added when needed if NO_3_^−^ was completely depleted during the week. Protein and DNA extractions were then performed from the biofilm (5^th^ transfer only) and the suspended biomass (all cultures).

### The Reference biofilm cultures (Ref300N-23C)

The carriers (Bioflow 9) containing the denitrifying biofilm (here named the Original Biofilm; OB) were taken from the denitrification reactor of the Montreal Biodome and frozen at −20°C in seawater with 20% glycerol (Laurin *et al.*, 2006) until use. Biomass from several carriers were thawed, scrapped, weighted and dispersed in the ASW at 0.08 g (wet weight)/mL. The biomass (5 mL/vial, 0.4 g of biofilm) was then distributed with a syringe and an 18_G_1½ needle into vials containing twenty free carriers and 60 mL ASW and supplemented with 300 mg NO_3_^−^-N/L (21.4 mM NO_3_^−^) and 0.15% (v/v) methanol (C/N=1.5). The vials were incubated at 23°C and shaken at 100 rpm (orbital shaker).

**Table 2.** Incubation conditions to measure the impact of specific parameters on the Reference biofilm cultures (Ref300N-23C)

| Tested conditions | $\text{NO}_3^-/\text{NO}_2^-$<br>mg-N/L(mM) | Methanol<br>% (v/v) | C/N | NaCl<br>% (w/v) | pH | Temp<br>°C |
| --- | --- | --- | --- | --- | --- | --- |
| $\text{NO}_3^-$ and methanol (C/N=1.5) | | | | | | |
| 300<sub>N-NO3</sub>-0.15%<sub>MeOH</sub>* | 300 (21.4) | 0.15 | 1.5 | 2.75 | 8.0 | 23 |
| 600<sub>N-NO3</sub>-0.3%<sub>MeOH</sub> | 600 (42.8) | 0.3 | 1.5 | 2.75 | 8.0 | 23 |
| 900<sub>N-NO3</sub>-0.45%<sub>MeOH</sub> | 900 (64.3) | 0.45 | 1.5 | 2.75 | 8.0 | 23 |
| 1500<sub>N-NO3</sub>-0.75%<sub>MeOH</sub> | 1500 (107) | 0.75 | 1.5 | 2.75 | 8.0 | 23 |
| 3000<sub>N-NO3</sub>-1.5%<sub>MeOH</sub> | 3000 (214) | 1.5 | 1.5 | 2.75 | 8.0 | 23 |
| $\text{NO}_2^-$ | | | | | | |
| 400<sub>N-NO2</sub> | 400 (28.6) | 0.15 | 1.5 | 2.75 | 8.0 | 23 |
| 200<sub>N-NO2</sub> | 200 (14.3) | 0.15 | 1.5 | 2.75 | 8.0 | 23 |
| 200<sub>N-NO3</sub>-200<sub>N-NO2</sub> | 200+200 (28.6) | 0.15 | 1.5 | 2.75 | 8.0 | 23 |
| Methanol (C/N variable) |  |  |  |  |  |  |
| 0%<sub>MeOH</sub> | 300 (21.4) | 0 | 0 | 2.75 | 8.0 | 23 |
| 0.05%<sub>MeOH</sub> | 300 (21.4) | 0.05 | 0.5 | 2.75 | 8.0 | 23 |
| 0.15%<sub>MeOH</sub>* | 300 (21.4) | 0.15 | 1.5 | 2.75 | 8.0 | 23 |
| 0.5%<sub>MeOH</sub> | 300 (21.4) | 0.5 | 5 | 2.75 | 8.0 | 23 |
| $\text{NO}_3^-$ (C/N variable) | | | | | | |
| 90<sub>N-NO3</sub> | 90 (6.4) | 0.15 | 5 | 2.75 | 8.0 | 23 |
| 300<sub>N-NO3</sub>* | 300 (21.4) | 0.15 | 1.5 | 2.75 | 8.0 | 23 |
| 900<sub>N-NO3</sub> | 900 (64.3) | 0.15 | 0.5 | 2.75 | 8.0 | 23 |
| 3000<sub>N-NO3</sub> | 3000 (214) | 0.15 | 0.15 | 5 | 8.0 | 23 |
| pH** |  |  |  |  |  |  |
| pH4 | 300 (21.4) | 0.15 | 1.5 | 2.75 | 4.0 | 23 |
| pH6 | 300 (21.4) | 0.15 | 1.5 | 2.75 | 6.0 | 23 |
| pH8* | 300 (21.4) | 0.15 | 1.5 | 2.75 | 8.0 | 23 |
| pH10 | 300 (21.4) | 0.15 | 1.5 | 2.75 | 10.0 | 23 |
| Temperature** |  |  |  |  |  |  |
| 5°C | 300 (21.4) | 0.15 | 1.5 | 2.75 | 8.0 | 5 |
| 15°C | 300 (21.4) | 0.15 | 1.5 | 2.75 | 8.0 | 15 |
| 23°C* | 300 (21.4) | 0.15 | 1.5 | 2.75 | 8.0 | 23 |
| 30°C | 300 (21.4) | 0.15 | 1.5 | 2.75 | 8.0 | 30 |
| 36°C | 300 (21.4) | 0.15 | 1.5 | 2.75 | 8.0 | 36 |
| NaCl** |  |  |  |  |  |  |
| 0% NaCl | 300 (21.4) | 0.15 | 1.5 | 0 | 8.0 | 23 |
| 1% NaCl | 300 (21.4) | 0.15 | 1.5 | 1.0 | 8.0 | 23 |
| 2.75%<sub>NaCl</sub>* | 300 (21.4) | 0.15 | 1.5 | 2.75 | 8.0 | 23 |
| 5% NaCl | 300 (21.4) | 0.15 | 1.5 | 5.0 | 8.0 | 23 |
| 8% NaCl | 300 (21.4) | 0.15 | 1.5 | 8.0 | 8.0 | 23 |

\*\*The Reference biofilm cultures were incubated in the prescribed conditions for two days, and then transferred in fresh medium.

**Figure 2.**
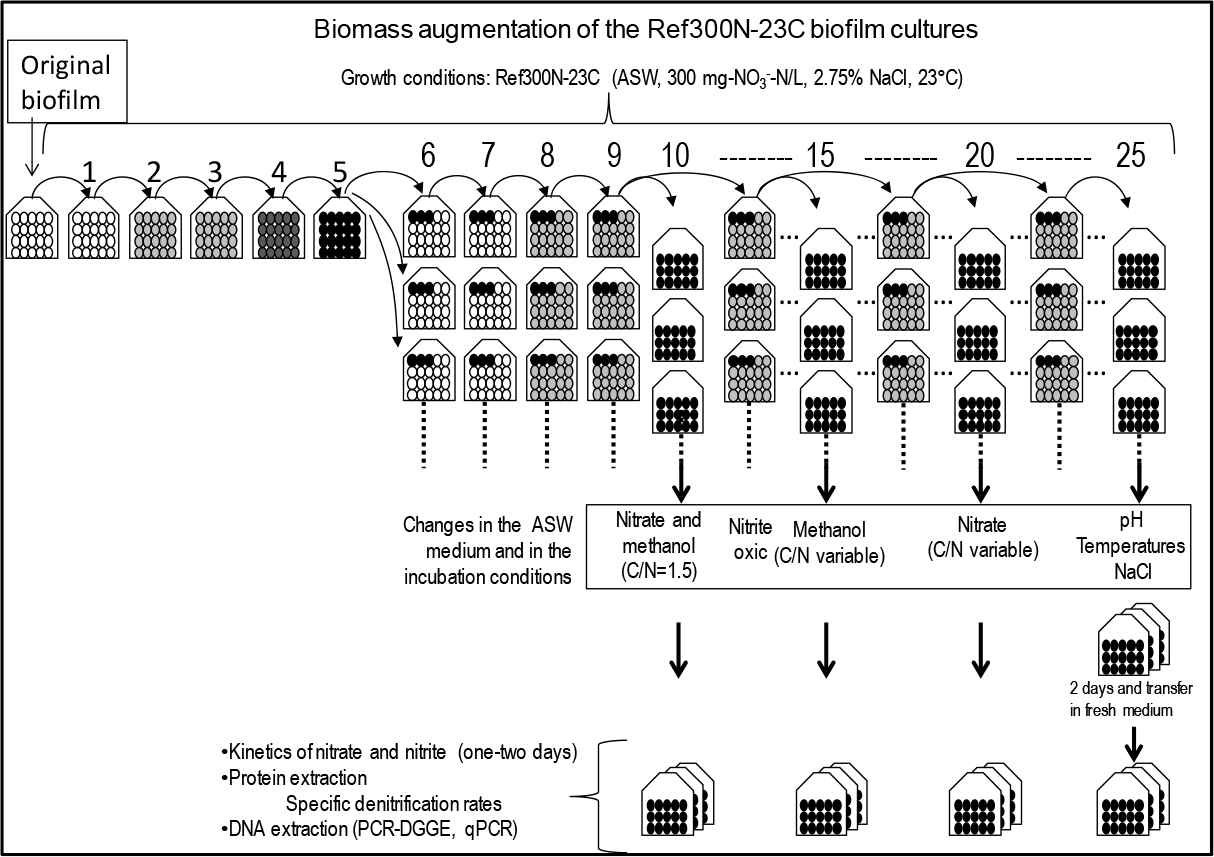
Schematic of the conditions used to test the denitrification capacity of the Ref300N-23C biofilm cultures. - The original biofilm was thawed, scrapped from the carriers and distributed to vials containing the ASW medium supplemented with 300 mg NaNO_3_-N/L and 0.15% methanol and 20 free carriers. These carriers were transferred 5 times in fresh medium. At the 5^th^, 10^th^, 15^th^ and 20^th^ carrier-transfers, series of three colonized carriers were distributed into other vials containing fresh culture medium and 17 new carriers. The vials were cultured and the carriers transferred as before.
- At the 10^th^, 12^th^, 15^th^ and 20^th^ carrier-transfers, series of 15 freshly colonized carriers were transferred into vials containing the prescribed medium to assess the impact in varying the NO_3_^−^, NO_2_^−^ and methanol concentrations or the oxic conditions on the denitrifying activities.
- At the 25^th^ transfer, series of 15 freshly colonized carriers were transferred into vials containing the prescribed medium to assess the impact in varying the pH, the temperature and the NaCl concentrations on the denitrifying activities. The biofilm was cultured for two days in these prescribed conditions before the carriers to be transferred in fresh prescribed medium.
- Kinetics of NO_3_^−^ and NO_2_^−^ were monitored at regular intervals. Protein and DNA extractions were then performed from the biofilm and the suspended biomass.

In average once a week, the twenty carriers were taken out of the vial, gently washed to remove the excess medium and the free bacteria, transferred into a new vial containing fresh anaerobic medium, and then incubated again under the same conditions (Fig. 1). Samples were taken each one or two days to measure the concentrations of NO_3_^−^ and NO_2_^−^ in the growth medium. The residual suspended biomass was taken for DNA extraction. Methanol and NaNO_3_ were added when needed if NO_3_^−^ was completely depleted during the week. During the fifth carrier-transfer cultures, multiple samples (500 μL) were collected at regular interval for 1-3 days to determine the NO_3_^−^ and NO_2_^−^ concentration. After this monitoring, total biomass in a whole vial was determined by protein quantification.

### Impact of physico-chemical changes on the Reference biofilm cultures

The carriers containing the Reference biofilm cultures were used as starting material to test the impact on the denitrification performance of the biofilm by varying specific physico-chemical parameters (Table 2). Figure 2 described in detail the protocol. To perform these assays, fifteen carriers from the Reference biofilm cultures were distributed into vials in the prescribed conditions described in Table 2. For the assays that was performed with different pHs and temperatures and different NaCl concentrations, the vials were cultured for two days for the biofilm to adapt to the new culture conditions, and then the carriers were transferred into their respective fresh medium (Fig. 2). NO_3_^−^ and NO_2_^−^ concentrations in the vials were then monitored by collecting 500 μL medium samples at regular intervals for 1 to 3 days. Biomass was taken at the end of this monitoring to extract DNA and for protein quantification. Most assays were performed in triplicates.

### Acclimation of the biofilms to different culture conditions

The OB was taken and processed as described in *The Reference biofilm cultures* subsection. The formulation of the ASW medium was adjusted with the prescribed NaCl, NO_3_^−^, NO_2_^−^ and methanol concentrations, and pHs, and the vials were incubated at prescribed temperatures (Table 3; Fig. 3). Acclimation of the OB under oxic conditions were performed as for the Reference biofilm cultures but in 250-mL Erlenmeyer flasks with agitation (Table 3). The OB acclimation to the IO medium was performed as for the Reference biofilm cultures (Table 3). All biofilm cultures were shaken at 100 rpm (orbital shaker), or 200 rpm for the oxic cultures. Five carrier-transfer cultures were performed, after which the NO_3_^−^, NO_2_^−^ and protein concentrations were determined as described in *The reference biofilm cultures* subsection.

**Table 3.** Culture conditions used for the acclimation of the original biofilm.

| Name | Medium | NO<sub>3</sub><sup>-</sup> / NO<sub>2</sub><sup>-</sup><br>mg-N/L<br>(mM) | Methanol<br>% (v/v) | NaCl<br>% (w/v) | Temp<br>°C |
| --- | --- | --- | --- | --- | --- |
| 300N-23C<sup>a</sup> | ASW | 300 (21.4) | 0.15 | 2.75 | 23 |
| Oxic<sup>b</sup> | ASW | 300 (21.4) | 0.15 | 2.75 | 23 |
| 300N-30C | ASW | 300 (21.4) | 0.15 | 2.75 | 30 |
| 900N-23C | ASW | 900 (64.3) | 0.45 | 2.75 | 23 |
| 900N-30C | ASW | 900 (64.3) | 0.45 | 2.75 | 30 |
| 0% NaCl | ASW | 300 (21.4) | 0.15 | 0 | 23 |
| 0.5% NaCl | ASW | 300 (21.4) | 0.15 | 0.5 | 23 |
| 1% NaCl | ASW | 300 (21.4) | 0.15 | 1.0 | 23 |
| 5% NaCl | ASW | 300 (21.4) | 0.15 | 5.0 | 23 |
| 8% NaCl | ASW | 300 (21.4) | 0.15 | 8.0 | 23 |
| 2.75-5% NaCl<sup>c</sup> | ASW | 300 (21.4) | 0.15 | 2.75/5 | 23 |
| IO | IO | 300 (21.4) | 0.15 | 3.0<sup>d</sup> | 23 |
| 200-200N | ASW | 200N-NO<sub>3</sub><sup>-</sup><br>200N-NO<sub>2</sub><sup>-</sup> (28.6) | 0.15 | 2.75 | 23 |

The original biofilm was cultured in triplicates in these conditions at pH 8.0. The carriers were transferred 5 times in fresh medium (8 times for the 200-200N biofilm cultures). In gray are changed parameters from the Ref300N-23C biofilm cultures. IO: Instant Ocean medium.
<sup>a</sup> Reference biofilm cultures.
<sup>b</sup> Acclimation was performed under oxic conditions.
<sup>c</sup> The Ref300N-23C biofilm cultures were further transferred three times in ASW composed of 5% NaCl.
<sup>d</sup> Based on the amount of Na<sup>+</sup> and Cl<sup>-</sup> in Table 1. The exact amount of NaCl added in the IO medium is not known. For comparison, the amount of Na<sup>+</sup> and Cl<sup>-</sup> in the ASW medium is 3.2%.

### Determination of the denitrification rates

Measurements of NO_3_^−^ and NO_2_^−^ concentrations in the biofilm cultures were performed in all assays using ion chromatography as described by Mauffrey *et al.* (2017). The total biomass in a vial was measured by collecting the suspended biomass and the biomass attached to the carriers (Fig. 1 to 3). The amount of protein in the biomass was determined by the Quick Start™ Bradford Protein Assay (BioRad, Mississauga, ON, Canada). The denitrification rates were calculated by the linear portion of the NO_3_^−^ plus NO_2_^−^ concentrations (NO_x_) over time for each replicate (mM h^−1^). The specific denitrification rates were reported as the denitrification rates divided by the quantity of biomass (mg protein) in a vial (mM h^−1^ mg-protein^−1^). Ammonium concentration was measured using the colorimetric method described by Mulvaney (1996).

### DNA-based analyses

The DNA was extracted from the suspended biomass and the biofilm that was scrapped from the carriers as previously described (Geoffroy *et al.*, 2018). The PCR amplification of the V3 region of the 16S ribosomal RNA (rRNA) genes for denaturating gradient gel electrophoresis (DGGE) experiments was performed as described (Lafortune *et al.*, 2009) with the 341f and 534r primers (Table 4). Total DNA extracted from the JAM1 and NL23 planktonic pure cultures was used to perform PCR amplifications of the same region of the 16S rRNA genes. The resulting strain-derived amplicons were co-migrated with the biofilm-derived amplicons on DGGE gels to identify the DNA fragments associated to strain JAM1 and strain NL23 in biofilm samples.

Quantitative PCR assays (SYBR green) to measure the concentrations of *M. nitratireducenticrescens* JAM1/GP59 (*narG1*), *M. nitratireducenticrescens* GP59 (*nirK*), *M. nitratireducenticrescens* JAM1 (*tagH*) and *H. nitrativorans* NL23 (*napA*) (Table 4) in the biofilm cultures were performed as previously described (Geoffroy *et al.*, 2018).

**Figure 3.**
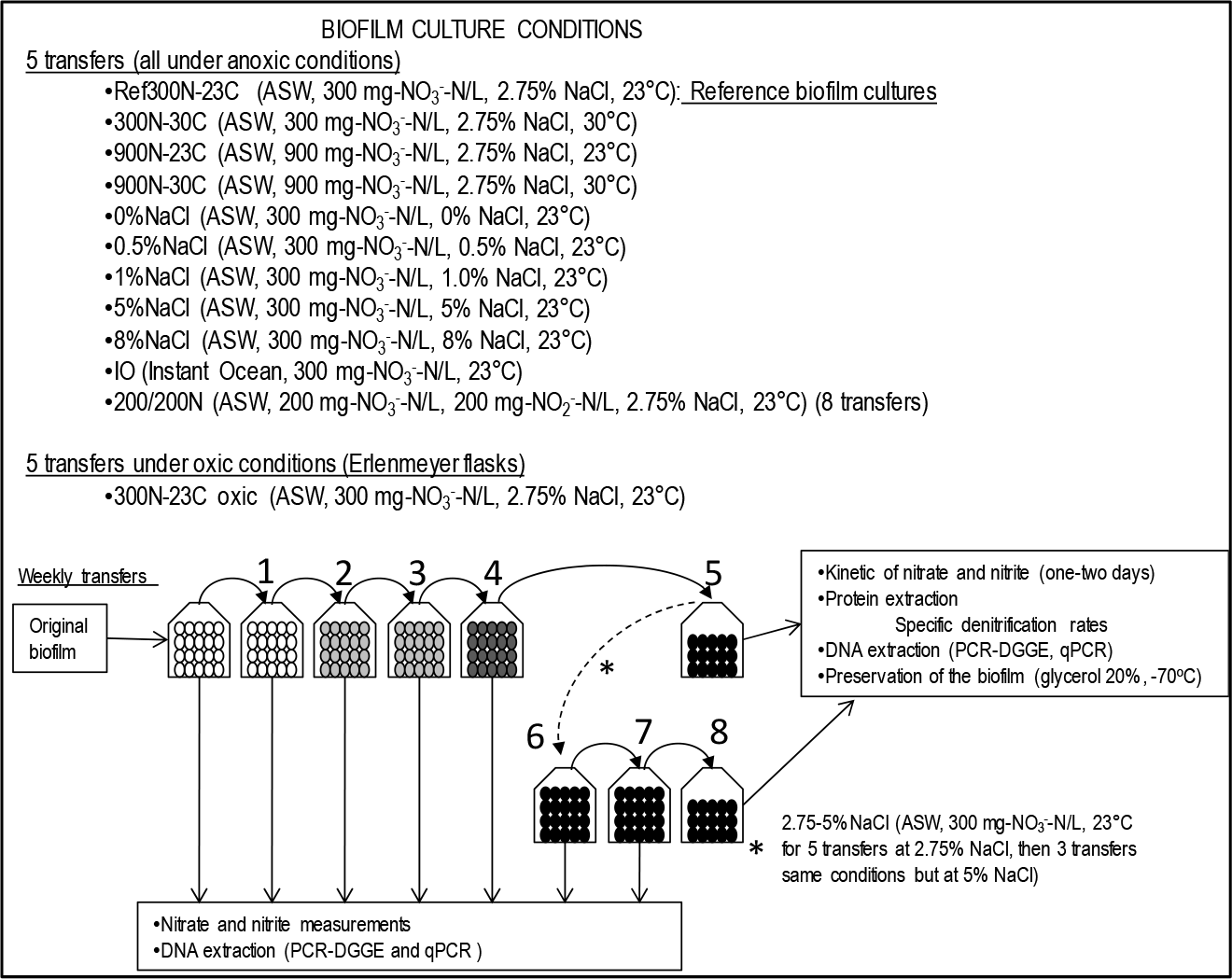
Schematic of the acclimation strategy used to derive the different biofilm cultures. The original biofilm was thawed, scrapped from the carriers and distributed to vials containing the ASW medium supplemented with prescribed concentrations of NO_3_^−^, NO_2_^−^, methanol and NaCl, or containing the IO medium (Table 2), and 20 free carriers. The vials were incubated at the prescribed temperatures and at pH 8 under anoxic or oxic conditions (Table 2). In average once a week, the carriers were transferred five times into fresh prescribed medium and incubated in the same conditions. For the 2.75-5%NaCl assay, the first five transfers were incubated in the same conditions of the Ref300N-23C biofilm cultures, then the three subsequent transfers were incubated in ASW medium with 5% NaCl. NO_3_^−^ and NO_2_^−^ concentrations were measured in each one or two days. Methanol and NaNO_3_ were added when needed if NO_3_^−^ was completely depleted during the week. DNA was extracted from suspended biomass for PCR-DGGE and qPCR assays. During the 5^th^ carrier-transfer cultures (or the 8^th^ for the 2.75-5%NaCl biofilm cultures), 15 carriers from each vial were transferred into their respective fresh medium, and concentrations of NO_3_^−^ and NO_2_^−^ were measured at regular intervals. Protein content in the whole vials was then measured.

### Statistics

Statistical significance in the denitrification rates between different types of cultures was performed with one-way ANOVA with Tukey's Multiple Comparison Test with the GraphPad Prism 5.0 software.

## Results

### Establishment of the Reference biofilm cultures (Ref300N-23C)

The Montreal Biodome operated a continuous fluidized methanol-fed denitrification system between 1998 and 2006. After the termination of the system, our group took the opportunity to preserve carriers with the denitrifying biofilm in glycerol at −20°C (Laurin *et al.*, 2006). The original biofilm (OB) from the frozen stock was scrapped from the carriers and the cells were dispersed. This dispersed biomass was used to inoculate vials containing new carriers to allow the development of a fresh denitrifying biofilm on the carrier surface. The vials were cultured under batch-mode anoxic conditions in the ASW medium (2.75% NaCl) with 300 mg NO_3_^−^-N/L (21.4 mM NO_3_^−^) and 0.15% (v/v) methanol (C/N=1.5) at 23°C. The choice of ASW medium was based on our ability to change the composition of the medium, as opposed to the commercial seawater medium (Instant Ocean) used by the Biodome. Each week for five weeks, only the carriers were transferred in fresh medium (Fig. 1). The biofilm cultured under these conditions is referred as the *Reference biofilm cultures* and named from here as the Ref300N-23C biofilm cultures (for 300 mg NO_3_^−^-N/L, 23°C). As its name implies, the conditions of these cultures were used as a reference to measure changes in the denitrifying activities resulting from changes in physico-chemical parameters in the culture medium.

**Table 4.**
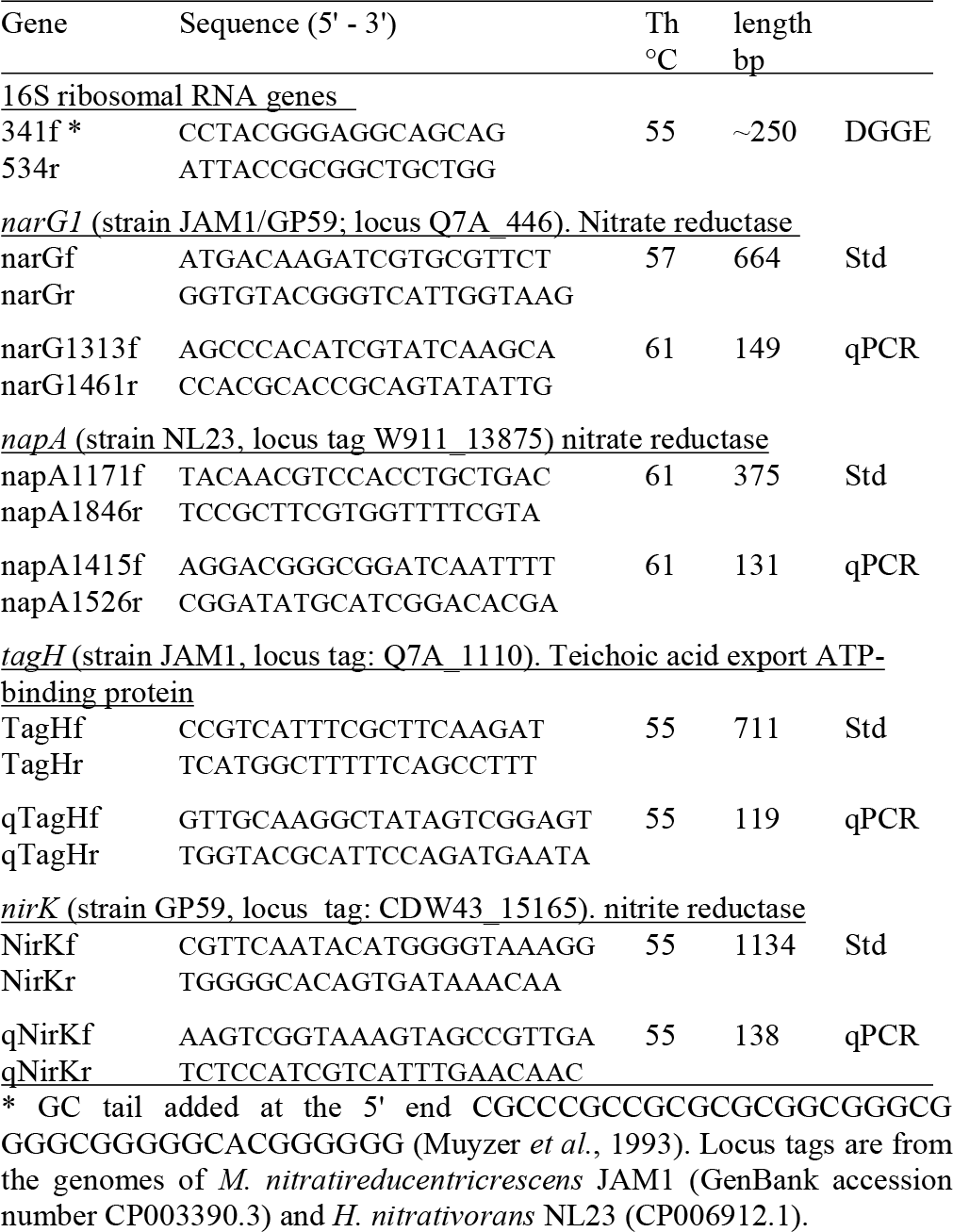
Primers used in this study.

The NO_3_^−^ and NO_2_^−^ concentrations were analyzed sporadically for the first four carrier-transfer cultures, and showed increasing rates of denitrifying activities with subsequent transfers. During the fifth carrier-transfer cultures, the concentrations of NO_3_^−^ and NO_2_^−^ were analyzed more thoroughly. NO_3_^−^ was consumed in 4 to 6 hours, with accumulation of NO_2_^−^ that peaked at 10 mM after 4 hours. NO_2_^−^ was completely consumed after 12 hours (Fig. 4A). The denitrification rates were calculated at 1.69 (±0.14) mM-NO_x_ h^−1^. Taking into account the biomass, the specific denitrification rates corresponded to 0.0935 (±0.0036) mM-NO_x_ h^−1^ mg-protein^−1^. The production of ammonia was negative in these conditions, which rules out dissimilatory NO_3_^−^ reduction to ammonium. Further carriers' transfers did not improve the specific denitrification rates.

The evolution of the bacterial community during its acclimation to these conditions was assessed by PCR-DGGE profiles for each carrier-transfer cultures (Fig. 4B). These profiles were similar after the third carrier-transfer cultures, suggesting stabilization of the bacterial community in these cultures. The most substantial changes in these profiles compared to the OB profile is the intensity of the DGGE band corresponding to strain JAM1 that was much stronger in all carrier-transfer cultures, whereas the DGGE band corresponding to strain NL23 was no longer visible after the second carrier-transfer cultures (Fig. 4B). qPCR analysis confirmed these results (Fig 4C). Strain JAM1 increased in concentration in the Ref300N-23C biofilm cultures from 5.6 × 10^3^ *narG1* copies/ng DNA in the OB to 2.0 × 10^5^ *narG1* copies/ng DNA in the fifth carrier-transfer cultures. Strain NL23 concentrations however dropped by three orders of magnitude from 8.7 × 10^4^ in the OB to 3.2 × 10^1^ *napA* copies/ng DNA in the fifth carrier-transfer cultures.

**Figure 4.**
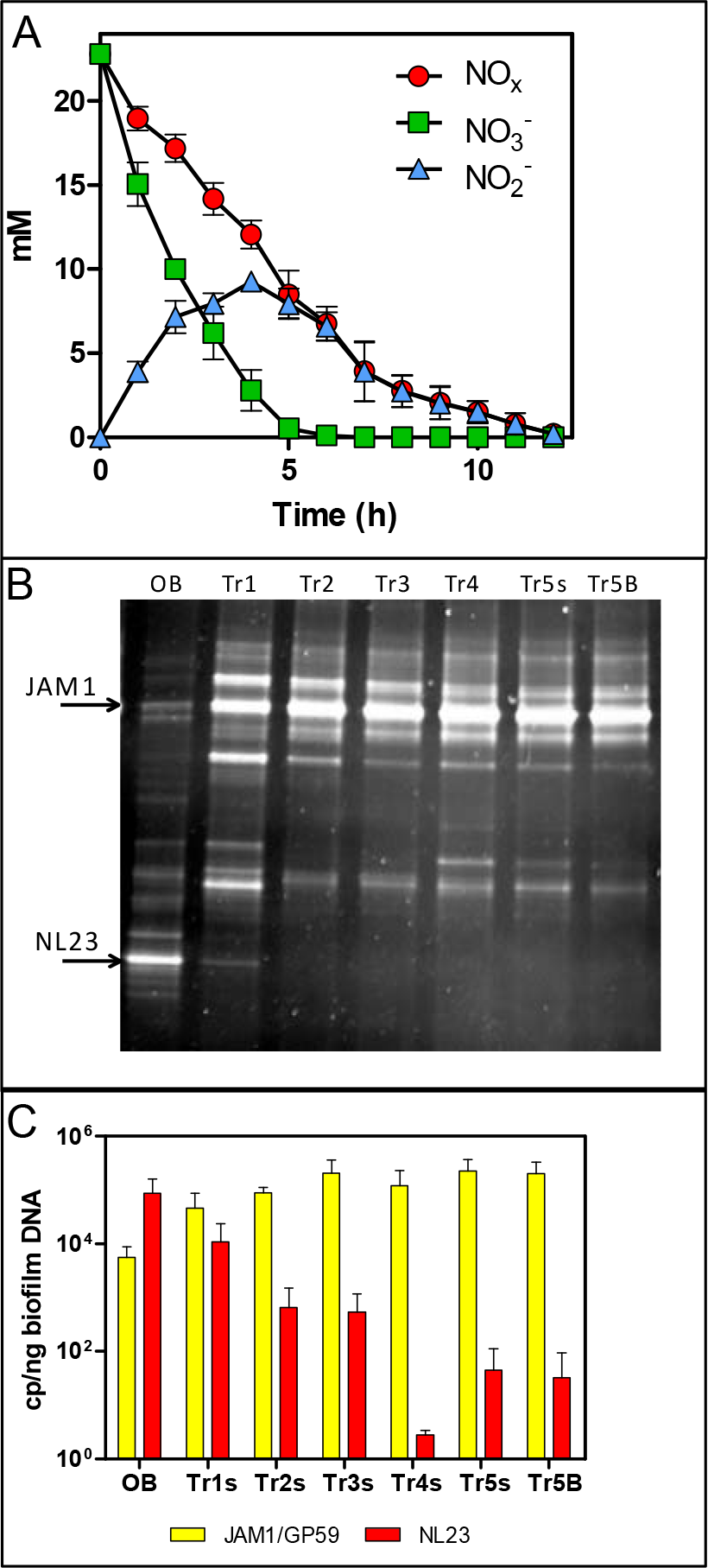
Dynamics of NO_3_^−^ and NO_2_^−^ concentrations and of the bacterial community in the Ref300N-23C biofilm cultures. **Panel A:** NO_3_^−^ and NO_2_^−^ concentrations were measured during the 5^th^ carriertransfer cultures. Results from triplicate biofilm cultures. **Panel B:** PCR-DGGE migration profiles representing the bacterial diversity during the first 5 carrier-transfer cultures. DNA extraction was performed on samples from the suspended biomass in the first 4 carrier-transfer cultures because the biofilm was not enough abundant on the carriers in these cultures. In the 5^th^ carrier-transfer cultures, samples were taken from the suspended biomass (Tr5s) and the biofilm (Tr5B). OB: Original biofilm. **Panel C.** Quantification of *Methylophaga nitratireducenticrescens* JAM1/GP59 (*narG1*) and *Hyphomicrobium nitrativorans* NL23 (*napA*) by qPCR in the 5 carrier-transfer cultures (Tr1 to Tr5). Results from 3 to 9 replicate cultures, of one to three different inoculums.

These results showed that important changes in the bacterial community occurred during the acclimation of the OB to the ASW medium under the batch-operating mode, with the important decrease in concentration of the main denitrifying bacterium in the biofilm, *H. nitrativorans* NL23, without the loss of denitrifying activities. We discovered later that a new *M. nitratireducenticrescens* strain, strain GP59 with full denitrification capacity, was enriched in these cultures (Geoffroy *et al.*, 2018). qPCR assays targeting *narG1* cannot discriminate strain JAM1 from strain GP59, as this gene is identical in nucleic sequences in both strains (Geoffroy *et al.*, 2018). Therefore, the concentration of strain JAM1 in the biomass derived from *narG1*-targeted qPCR assays and illustrated in Figure 4C reflects in fact the concentrations of both strains.

### Impact of physico-chemical changes on the Ref300N-23C biofilm

The colonized carriers from the Ref300N-23C biofilm cultures were exposed for a short period (few hours to few days) to a range of methanol, NO_3_^−^, NO_2_^−^ and NaCl concentrations, and to different pHs and temperatures (Table 2, Fig. 2). These conditions were chosen to assess the capacity range of the denitrifying activities of the biofilm (methanol and NO_3_^−^ concentrations), but also its resilience to adverse conditions that could occur during the operation of a bioprocess (*e.g.* abrupt change of pH, temperature or salt concentration).

Increasing the concentrations of NO_3_^−^ and methanol (fixed C/N at 1.5) showed increases in the denitrification rates compared to the original conditions of the Ref300N-23C biofilm cultures (300 mg-NO_3_^−^-N/L with 0.15% methanol), with the highest increases observed at 600 mg-NO_3_^−^-N/L with 0.3% methanol (58% increase), and at 900 mg-NO_3_^−^-N/L with 0.45% methanol (56% increase) (Fig. 5). Inhibition of the denitrifying activities occurred at 3000 mg-NO_3_^−^-N/L. This inhibition could have been caused by the methanol concentration (1.5%) in the medium. Variations in methanol concentrations (fixed concentration of NO_3_^−^ at 300 mg-NO_3_^−^-N/L) showed no improvement of the denitrification activities compared to the original conditions (Fig. 5). Absence of methanol showed very weak denitrifying activities as expected. Variations in NO_3_^−^ concentrations (fixed concentration of methanol at 0.15%) showed the highest denitrification rates (74% increase) at 900 mg-NO_3_^−^-N/L (Fig. 5). Denitrifying activities were observed this time at 3000 mg-NO_3_^−^-N/L. However, NO_3_^−^ consumption stopped after 5 days and 50% consumption. Addition of methanol (0.15%) allowed activities to resume for one day with a further 20% NO_3_^−^ consumption.

Exposure of the Ref300N-23C biofilm cultures to pH 4 and 6 showed 2.7 and 2.6-fold increases of the denitrification rates, respectively, compared to the original conditions (pH 8) (Fig. 5). However, the pH at the end of the assays (10 to 12 hours) increased at around 8 in the medium. At pH 10, the denitrifying activities were completely inhibited. For the temperature assays, the highest denitrification rates were recorded at 30 and 36°C (Fig. 5), with 53% and 62% increases, respectively, compared to the original conditions (23°C). At 5°C, the denitrifying activities were still occurring but at 4-time lower rates than at 23°C. At 15°C, the denitrification rates were not significantly different from the original conditions. Exposure of the Ref300N-23C biofilm concentrations ranging from 0% to 8% showed little effect on the denitrification rates (Fig. 5). Finally, presence of NO_2_^−^ has a strong effect on the denitrification rates with 11-fold and 6.6-fold decreases with 400 mg-NO_2_^−^-N/L and 200 mg-NO_2_^−^-N/L, respectively in the medium, compared to the original conditions (Fig. 5). Mixture of 200 mg-NO_2_^−^-N/L and 200 mg-NO_3_^−^-N/L in the medium had a less pronounce effect with a 3-fold decrease of the denitrification rates.

**Figure 5.**
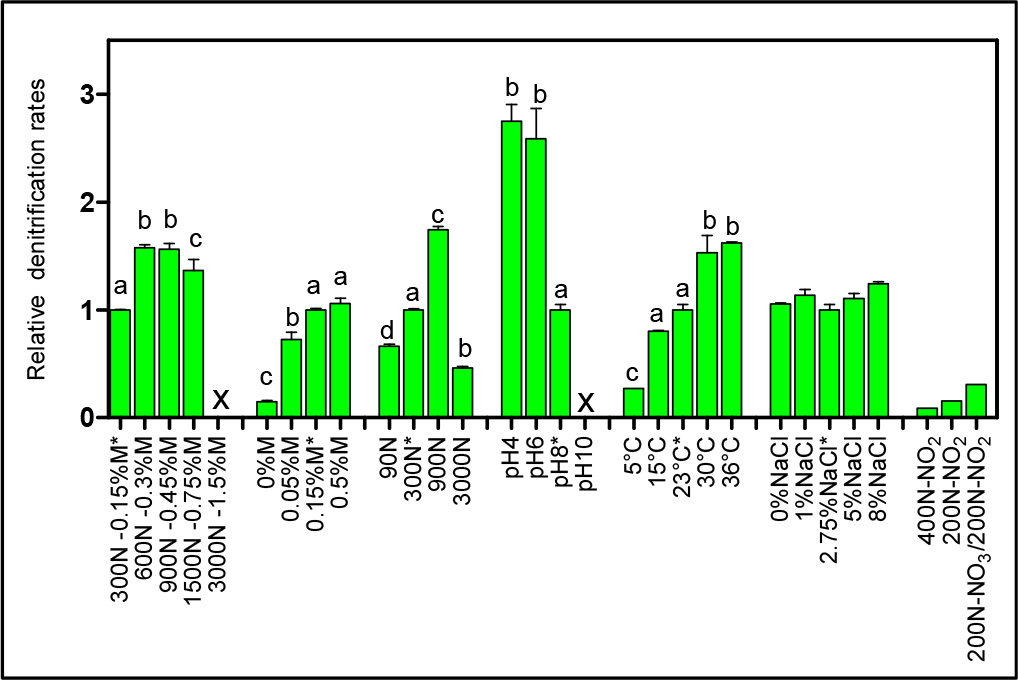
Denitrifying activities of the Ref300N-23C biofilm cultures impacted by different physico-chemical parameters. The Ref300N-23C biofilm cultures were exposed for few hours or few days to ASW-modified medium as described in Table 1. Denitrification rates are expressed relative to the denitrification rates of the original conditions of the Ref300N-23C biofilm cultures (set to one) identified by *. Significance between the denitrification rates were determined by one-way ANOVA. Rates with the same letter are not significantly different. X: no activities recorded. Average of triplicates cultures except for the 400N-NO_2_, 200N-NO_2_ and 200N-NO_3_/200N-NO_2_ conditions with one replicate. N: Nitrogen NO_3_^−^. M: methanol.

No significant differences in the quantity of biomass (protein amount per vial) was observed at the end of all these assays between vials. In all the tested conditions, the bacterial diversity profiles (PCR-DGGE profiles) showed no changes (Fig. 4B, lane Tr5B). The levels of *M. nitratireducenticrescens* JAM1/GP59 was around 10^5^ *narG1* copies/ng DNA and of strain NL23 between 10^1^ and 10^−1^ *napA* copies/ng DNA.

### Acclimation of the original biofilm to optimal NO_3_^−^ concentrations and temperatures

As described above, the Ref300N-23C biofilm cultures exposed for a short period to 900 mg-NO_3_^−^-N/L or to 300 mg-NO_3_^−^-N/L at 30 and 36°C showed between 53% to 74% increases in the denitrification rates compared to the original conditions (Fig. 5). Based on these results, we acclimated the OB under optimal cultures conditions in the attempt to derive a biofilm with higher denitrifying capacity. As before, the dispersed biomass of the OB was used as inoculums to colonize new carriers. Five carrier-transfer cultures were carried out in ASW medium supplemented with 900 mg-NO_3_^−^-N/L, and incubated at either 23°C or 30°C (here named 900N-23C and 900N-30C, respectively), or at 300 mg-NO_3_^−^-N/L at 30°C (here named 300N-30C). In parallel, the Ref300N-23C biofilm cultures were derived by the same protocol (Table 3, Fig. 3). Compared to the Ref300N-23C biofilm cultures, the denitrification rates of the 900N-30C biofilm cultures increased by 61% (Fig. 6A). The two other biofilm cultures showed both 32% increases in these rates. The amounts of biomass per vials were not highly significantly different between the four cultures (Fig. 6A).

The bacterial diversity profiles (derived by PCR-DGGE) of these four acclimated biofilm cultures were all similar (same profile illustrated in Fig. 4B, lane Tr5B). qPCR analyses showed the same trend in the dynamics of *M. nitratireducenticrescens* JAM1/GP59 and *H. nitrativorans* NL23 populations. *M. nitratireducenticrescens* JAM1/GP59 concentrations increased by two orders of magnitude (1.9 to 2.3 × 10^5^ *narG1* copies/ng DNA), and strain NL23 concentrations decreased by 2-3 orders of magnitude (6.1 to 18 × 10^1^ *napA* copies/ng DNA) in the four biofilm cultures compared to the OB (Fig. 6B). After the isolation of strain GP59 and the sequencing of its genome, qPCR primers specific to strain JAM1 and to strain GP59 were developed (Geoffroy *et al.*, 2018). The concentrations of these two strains were then measured in the OB and in the four biofilm cultures. Between 1.6 and 2.9 × 10^5^ *nirK* copies/ng DNA of strain GP59 were measured in these biofilm cultures, which is three orders of magnitude higher than what was measured in the OB (1.6 × 10^2^ *nirK* copies/ng DNA). The concentrations of strain JAM1 in the four biofilm cultures ranged from 2.4 to 8.5 ×10^2^ *tagH* copies/ng DNA, which is 3 to 10 times lower than its concentration in the OB (2.5 ×10^3^ *tagH* copies/ng DNA) (Fig. 6B).

### Acclimation of the original biofilm to different NaCl concentrations

In the first set of assays, we showed that the Ref300N-23C biofilm cultures could sustain high variations of NaCl concentrations for a short period without affecting the denitrification capacity. Based on these results, we tested the capacity of the OB to acclimate to a range of NaCl concentrations. The dispersed biomass of the OB was cultured during five carrier-transfers under anoxic conditions in ASW-modified medium with NaCl concentrations ranging from 0% to 8% (Table 3, Fig. 3). The ASW used for the Ref300N-23C biofilm cultures contains 2.75% NaCl.

In biofilm cultures acclimated to low NaCl concentrations (0, 0.5 and 1.0%), the amounts of biomass developed were 1.3 to 2 times lower than in the Ref300N-23C biofilm cultures (Fig. 6A). This lower development did not impair very much the denitrification rates of the 0%NaCl and 0.5%NaCl biofilm cultures, but a 2.2-fold decrease of these rates in the 1%NaCl biofilm cultures was observed compared to the Ref300N-23C biofilm cultures (Fig. 6A). The 5%NaCl and 8%NaCl biofilm cultures were strongly affected by poor biofilm development on the carriers, which was 50-100 less abundant in biomass than in the Ref300N-23C biofilm cultures (Fig. 6A). In these biofilm cultures, NO_3_^−^ took more than 24 hours to be consumed and NO_2_^−^ consumption was minimal. This result contradicts what was obtained in the first set of assays where the Ref300N-23C biofilm cultures subjected for 3 days to 5% and 8% NaCl did not affect the denitrification performance. To investigate this further, another Ref300N-23C biofilm cultures were derived, but after the fifth transfer cultures, the carriers were transferred in 5% NaCl ASW medium (here named 2.75-5%NaCl) and cultured for another three carriers' transfers (around 20 days). These biofilm cultures were able to sustain this high salt concentration. Compared to the Ref300N-23C biofilm cultures, the 2.75-5%NaCl biofilm cultures had a 74% increase in the denitrification rates and the biomass was twice the amount (Fig. 6A).

**Figure 6.**
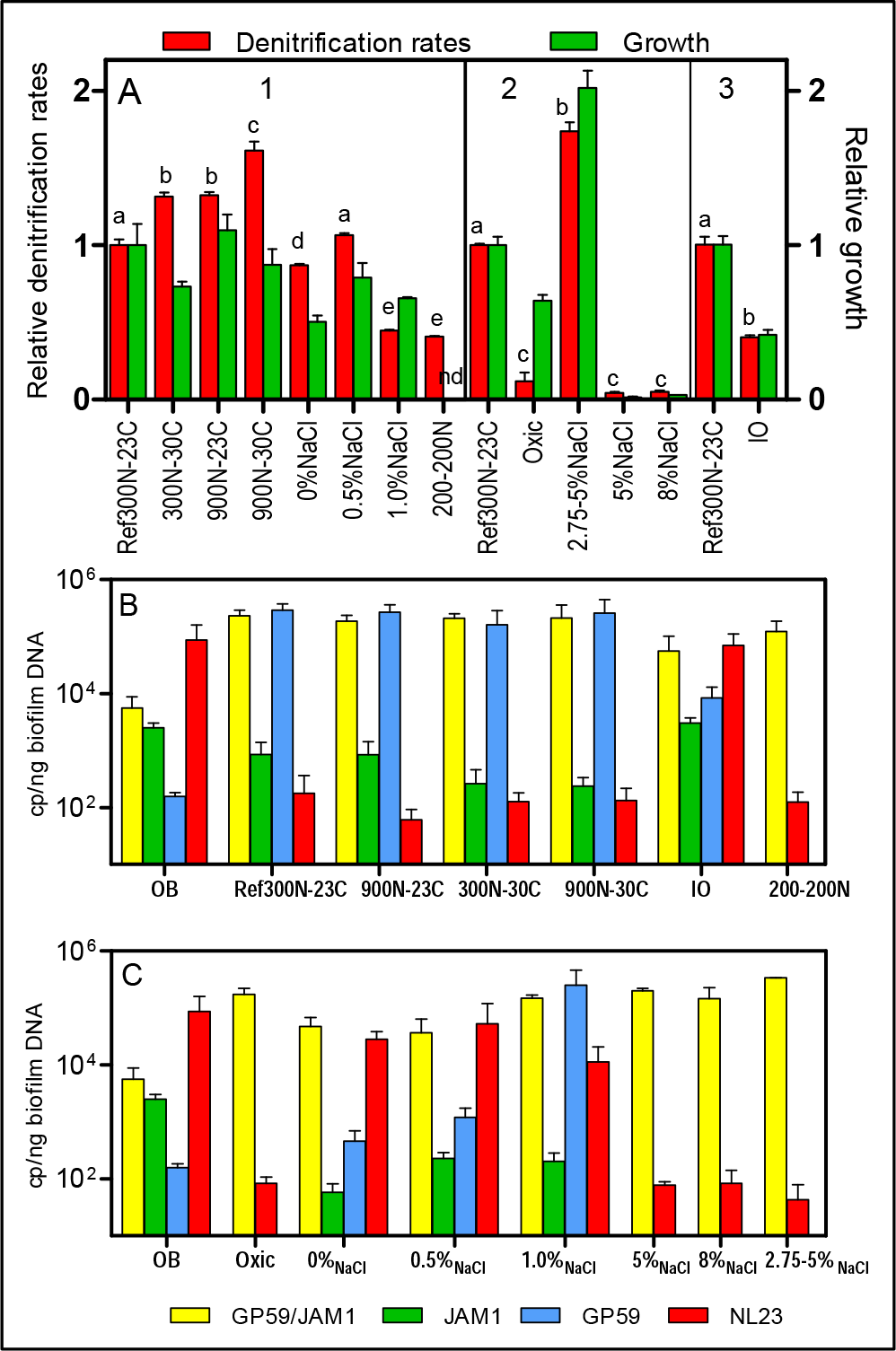
Denitrifying activities and dynamics of *H. nitrativorans* NL23 and *M. nitratireducenticrescens* populations in the biofilm cultures. **Panel A.** Three sets of cultures assays (1, 2, 3) were performed with the Ref300N-23C biofilm cultures in each set for comparison. Denitrification rates (mM h^−1^ vial^−1^) are expressed relative to the denitrification rates of the Ref300N-23C biofilm cultures set to one. Growth (mg protein vial^−1^) are expressed relative to the growth of the Ref300N-23C biofilm cultures. Significance between the denitrification rates were determined by one-way ANOVA or unpaired T-test within the same set of cultures assays. Rates with the same letter are not significantly different. See Table 2 and Material & Methods for nomenclature and culture conditions. Average of triplicates cultures. nd: not done. **Panels B and C.** Quantification of *M. nitratireducenticrescens* JAM1/GP59 (*narG1*), *M. nitratireducenticrescens* JAM1 (*tagH*), *M. nitratire-ducenticrescens* GP59 (*nirK*) and *Hyphomicrobium nitrativorans* NL23 (*napA*) by qPCR in the corresponding biofilm cultures. Results from triplicate cultures. OB: Original biofilm.

The PCR-DGGE migrating profiles showed the persistence of strain NL23 in the 0%NaCl and 0.5%NaCl biofilm cultures. In the 1%NaCl biofilm cultures, however, strain NL23 was barely visible in DGGE profiles (not shown). Persistence of strain NL23 was confirmed by qPCR where its concentrations in the 0%NaCl, 0.5%NaCl and 1.0%NaCl biofilm cultures (1.1 to 5.3 × 10^4^ *napA* cp/ng DNA) were at the same order of magnitude than in the OB (8.7 × 10^4^ *napA* cp/ng DNA) (Fig. 6C). As observed before, strain NL23 concentrations decreased by two-three orders of magnitude in the Ref300N-23C, 5%NaCl, 8%NaCl and 2.75-5%NaCl biofilm cultures (4.3 to 18 × 10^1^ *napA* cp/ng DNA) compared to the OB (Fig. 6C). The levels of *M. nitratireducenticrescens* JAM1/GP59 increased by one order of magnitude in the 0%NaCl and 0.5%NaCl biofilm cultures (*ca.* 4 × 10^4^ *narG1* cp/ng DNA) compared to the OB (5.6 × 10^3^ *narG1* cp/ng DNA), at similar levels than strain NL23 (Fig. 6C). However, in the 1%NaCl biofilm cultures, the concentrations of *M. nitratireducenticrescens* JAM1/GP59 increased by another order of magnitude (1.5 × 10^5^ *narG1* cp/ng DNA), which is similar than that found in the Ref300N-23C biofilm cultures (2.3 × 10^5^ *narG1* cp/ng DNA) (Fig. 6B and 6C). As observed before, the concentrations of strain JAM1 decreased by approximately one order of magnitude in these cultures (6 to 23 × 10^2^ *tagH* cp/ng DNA) when compared to the OB (2.5 × 10^3^ *tagH* cp/ng DNA) (Fig. 6C). The concentrations of strain GP59 in the 0% NaCl biofilm cultures (4.6 × 10^2^ *nirK* cp/ng DNA) were at similar level than in the OB (1.6 × 10^2^ *nirK* cp/ng DNA). It increased by one order of magnitude in the 0.5%NaCl biofilm cultures (1.2 × 10^3^ *nirK* cp/ng DNA) and by two orders of magnitude in the 1.0%NaCl biofilm cultures (2.5 × 10^5^ *nirK* cp/ng DNA), similar to the levels reached in the Ref300N-23C biofilm cultures (2.9 × 10^5^ *nirK* cp/ng DNA) (Fig. 6B and 6C). All these results showed that strain GP59 was favored in the ASW medium with increasing concentrations of NaCl, which was the opposite for strain NL23 in these cultures.

### Acclimation of the original biofilm: ASW medium vs IO medium

The important decrease of strain NL23 and the enrichment of strain GP59 in the Ref300N-23C biofilm cultures suggest that the switch from the continuous-operating mode that prevailed in the Biodome reactor to the batch mode in our biofilm cultures could have generated these specific changes in the *H. nitrativorans* and *M. nitratireducenticrescens* populations. Another factor that could have influenced these changes is the ASW medium that we used instead of the
Instant Ocean (IO) from Aquatic systems Inc. used by the Biodome (Table 1). The dispersed cells of the OB were therefore acclimated to the IO medium, under the same conditions than the Ref300N-23C biofilm cultures (performed in parallel). The NO_3_^−^ and NO_2_^−^ dynamics in the Ref300N-23C biofilm cultures were similar than reported in Figure 4A. The denitrification rates were calculated at 1.89 ± 0.10 mM-NO_x_ h^−1^, and the specific denitrification rates corresponded to 0.0684 ±0.0005 mM-NO_x_ h^−1^ mg-protein^−1^. In the IO biofilm cultures, the denitrification rates (0.76 ± 0.03 mM-NO_x_ h^−1^) were 2.5 lower than in the Ref300N-23C biofilm cultures (Fig. 6A). However, because less biomass developed in the IO biofilm cultures (Fig. 6A), its specific denitrification rates (0.0661 ±0.0063 mM-NO_x_ h^−1^ mg-protein^−1^) were the same than in the Ref300N-23C biofilm cultures.

As before, PCR-DGGE migration profiles and qPCR assays showed the important decrease in strain NL23 concentrations during the five carriers' transfers in the Ref300N-23C biofilm cultures. Surprisingly, the DGGE band corresponding to strain NL23 was still present in the IO biofilm cultures (not shown). This result was confirmed by qPCR assays where the concentrations of strain NL23 in the IO biofilm cultures (Fig. 6B) were at similar levels (7.0 × 10^4^ *napA* cp/ng DNA) than in the OB (8.7 × 10^4^ *napA* copies/ng DNA). In the IO biofilm cultures, the concentrations of strain JAM1 and strain GP59 were both approximately one order of magnitude lower than those of strain NL23 (Fig. 6B).

### Acclimation of the original biofilm to oxic conditions and to NO_2_^−^

The dispersed cells of the OB were cultured under the same conditions of the Ref300N-23C biofilm cultures except that the cultures were incubated under oxic conditions. Growth was not impaired in these cultures, but the denitrification rates were 8.3 times lower than in the Ref300N-23C anoxic biofilm cultures (Fig. 6A). The OB acclimation to a mix of 200 mg-NO_3_^−^-N/L and 200 mg-NO_2_^−^-N/L (here named 200-200N) generated biofilm cultures with denitrification rates 2.5-times lower than those in the Ref300N-23C biofilm cultures (Fig. 6A). The proportions of *M. nitratireducenticrescens* JAM1/GP50 and strain NL23 in these two types of cultures were similar than in the Ref300N-23C biofilm cultures (Fig. 6B and 6B).

## Discussion

The acclimation of the OB in the ASW medium by colonization of new carriers and their successive transfers in fresh medium had a strong impact on the concentration of *H. nitrativorans* NL23. From the second carriers’ transfer, a decrease of one order of magnitude was noticed in the levels of strain NL23, and resulted in further decreases after several transfers. However, these important decreases did not translate into impaired denitrification rates. On the opposite, the concentrations of *M. nitratireducenticrescens* increased by two orders of magnitude in the biofilm cultures after the third carriers’ transfer. We already reported in Geoffroy *et al.* (2018) that *H. nitrativorans* NL23 was displaced in the Ref300N-23C biofilm cultures by a new strain of *M. nitratireducenticrescens*, strain GP59, that can perform complete denitrification. The increase of *M. nitratireducenticrescens* concentrations involved mainly strain GP59, which was at a very low level in the OB, but increased by three order of magnitude in ASW-derived biofilm cultures. Strain JAM1 strayed approximately at the same levels in the OB and the Ref300N-23C biofilm cultures. The reasons why strain GP59 did not appear in first instance in the denitrification system of the Biodome are obscure. This may be related to the continuous mode operation used in the Biodome system versus the batch mode cultures used in our studies.

Measurements of the denitrifying activities immediately after transferring the carriers in ASW-modified media allowed the assessment of the capacity of the Ref300N-23C biofilm cultures. These biofilm cultures can sustain denitrifying activities in most of the tested conditions. Denitrifying activities were inhibited when the biofilm cultures were exposed at pH 10 or with 1.5% methanol, and were strongly impacted by the presence of NO_2_^−^. Low pHs had a strong effect on the denitrification rates, with a 2.7-fold increase at pH 4.0 than at pH 8.0 in the original conditions. The ASW medium is slightly turbid at pH 8.0, but was clear at pH 4.0 and pH 6.0. This result suggests better solubility of elements at these pHs, which may have influenced the dynamics of NO_3_^−^ processing by the bacterial community. At the end of the assays (10-12 hours), however, the pH in the medium increased to reach pH 8.0. Denitrification is an alkalization process that leads to the increase of pH in the medium (Lee and Rittmann, 2003). Phosphate used as buffering system explains the stabilization of the pH at around 8.0.

In the second set of assays, a series of biofilm cultures were performed, in which the bacterial community of the OB was acclimated to different conditions for five weeks. The best denitrification performance occurred in the 900N-30C biofilm cultures, which allowed a 61% increase in the denitrification rates compared to the Ref300N-23C biofilm cultures. We did not try low pHs in our acclimation cultures because of the rapid fluctuations of the pH during the first set of assays. The NaCl concentration was a critical factor for the persistence of *H. nitrativorans* NL23 in the biofilm cultures. In the ASW-modified medium with low NaCl concentrations (0%, 0.5% and 1.0%), high levels of *H. nitrativorans* NL23 were found. Martineau *et al.* (2015) showed that strain NL23 can sustain good growth and denitrifying activities up to 1% NaCl in planktonic pure cultures, but underwent substantial decrease in these features at 2% NaCl. In contrast, growth of *Methylophaga* spp. requires Na^+^ (Bowman, 2005). In the 0%NaCl biofilm cultures, low concentration of Na^+^ (0.05%; originating from NaNO_3_ and Na_2_HPO_4_, Table S1) may have impaired the growth of *M. nitratireducenticrescens* JAM1 and GP59. Consequently, strain *H. nitrativorans* NL23 may have better competed against the *Methylophaga* strains in these cultures. On the opposite, the rapid processing of NO_3_^−^ by *M. nitratireducenticrescens* (Mauffrey *et al.*, 2015) in the Ref300N-23C biofilm cultures (containing 2.75% NaCl) with transient accumulation of NO_2_^−^ may have impaired the growth of *H. nitrativorans* NL23. The adverse environment for strain NL23 would have favored the immergence of strain GP59 that took over strain NL23 in completing the full denitrification pathway.

An interesting result was to find the persistence of strain NL23 in the biofilm cultivated under the same conditions than the Ref300N-23C biofilm cultures but with the commercial medium (IO) that was used in the Biodome denitrification system. The concentrations of *M. nitratireducenticrescens* JAM1/GP59 and *H. nitrativorans* NL23 were similar in the IO biofilm cultures. Although the IO biofilm cultures generated less biomass than the Ref300N-23C biofilm cultures, the specific denitrification rates were the same between the two types of cultures, suggesting similar dynamics of NO_3_^−^ processing by both bacterial communities. The reason of this fundamental difference has to be found in the composition of the seawater medium. Based on limited information on the composition of the IO medium, a difference that might impair the growth of strain NL23 is the lack of bicarbonate and carbonate in the ASW medium. In the IO medium, these elements are required for buffering the medium, which is performed by phosphate in the ASW medium. *Hyphomicrobium* spp. use the serine pathway for the C1 assimilation, which requires one molecule of CO_2_ and two molecules of formaldehyde in each cycle forming a three-carbon intermediate, while in *Methylophaga* spp. the ribulose monophosphate cycle requires three molecules of formaldehyde (Anthony, 1982). In our anoxic cultures, which were a closed environment, the ambient atmospheric CO_2_ was probably rapidly consumed.

The co-occurrence of *Methylophaga* spp. and *Hyphomicrobium* spp. observed in our studies has also been reported in two other denitrifications systems. Osaka et al. (2008) studied a laboratory-scale continuously stirred tank reactor with synthetic wastewater and sludge adapted to denitrification conditions. Among the tested assays, the reactor was fed with methanol and acclimated with increasing concentrations of NaCl. This reactor could perform high levels of NO_3_^−^ removal at up to 3% NaCl, but a drastic decrease in the removal efficiency occurred at 4% NaCl. Further increase in NaCl concentrations failed to get denitrifying activities in the reactor. Co-occurrence of *Methylophaga* spp. and *Hyphomicrobium* spp. was revealed in cloned 16S rRNA gene libraries derived from the 0% and 4% NaCl synthetic wastewater media, with respectively 35% and 20% of the 16S clones affiliated to *Hyphomicrobium* spp., and 8% and 11% affiliated to *Methylophaga* spp. The other denitrification system is similar to the one that was used at the Biodome, which is a methanol-fed fluidized-bed continuous denitrification system treating a marine aquarium in Helsinki, Finland (Rissanen *et al.*, 2016). In this system, the biofilm developed on oolitic sand. *Methylophaga* spp. and *Hyphomicrobium* spp. were found in high proportions although high variations occurred between samples taken at different years.

## Conclusions

This study shows the plasticity of the denitrifying biofilm to adapt to different conditions. The biofilm can sustain denitrifying activities in most of the tested conditions. We also showed an important evolution of the *M. nitratireducenticrescens* population with the emergence of a new denitrifying strain belonging to this species. *H. nitrativorans* NL23 was the most affected by these changes where the NaCl concentration and the type of seawater medium were factors influencing its persistence or its disappearance. A third part of this study is reported by Villemur *et al.* (2019), where we have looked at the overall microbial community of our biofilm cultures by using culture-dependent, 16S metagenomic and metatranscriptomic approaches aiming to assess the contribution of other denitrifying bacteria in the biofilm cultures.

## ACKNOWLEDGEMENTS

We thank Karla Vasquez for her technical assistance.

## Funding

This research was supported by a grant to Richard Villemur from the Natural Sciences and Engineering Research Council of Canada # RGPIN-2016-06061.

